# Flooding and hydrologic connectivity modulate community assembly in a dynamic river-floodplain ecosystem

**DOI:** 10.1101/557405

**Authors:** Stefano Larsen, Ute Karaus, Cecile Claret, Ferdinand Sporka, Ladislav Hamerlík, Klement Tockner

## Abstract

Braided river floodplains are highly dynamic ecosystems, where aquatic communities are strongly regulated by the hydrologic regime. So far, however, understanding of how flow variation influences assembly mechanisms remains limited.

We collected benthic chironomids and oligochaetes over a year across a lateral connectivity gradient in the semi-natural Tagliamento River (Italy). Four bankfull flood events occurred during the study, allowing the assessment of how flooding and hydrologic connectivity mediate the balance between stochastic and deterministic community assembly.

While invertebrate density and richness were positively correlated with connectivity, diversity patterns showed no significant correlation. Species turnover through time increased with decreasing connectivity. Contrary to expectations, hydrologic connectivity did not influence the response of community metrics (e.g. diversity, density) to floods.

Invertebrate composition was weakly related to connectivity, but changed predictably in response to floods. Multivariate ordinations showed that faunal composition diverged across the waterbodies during stable periods, reflecting differential species sorting across the lateral gradient, but converged again after floods. Stable hydrological periods allowed communities to assemble deterministically with prevalence of non-random beta-diversity and cooccurrence patterns and larger proportion of compositional variation explained by local abiotic features. These signals of deterministic processes clearly declined after flooding events. This occurred despite no apparent evidence of flood-induced homogenisation of habitat conditions.

This study is among the first to examine the annual dynamic of aquatic assemblages across a hydrologic connectivity gradient in a natural floodplain. Results highlight how biodiversity can exhibit complex relations with hydrologic connectivity. However, appraisal of the assembly mechanisms through time indicated that flooding shifted the balance from deterministic species sorting across floodplain habitats, towards stochastic processes related to organisms redistribution and the likely resetting of assembly to earlier stages.

## Introduction

Braided rivers floodplains are among the most dynamic ecosystems, constituting a shifting mosaic of habitat patches with high spatio-temporal turnover rates [1,2]. The complex interaction between floodplain topography and variation in river flow and sediment transport maintains a distinct gradient of lateral hydrologic connectivity, which facilitates the coexistence of numerous aquatic, amphibian, and terrestrial species [2]. This connectivity is defined as the permanent or temporary links between the main stem of the river and the diverse waterbodies across the alluvial floodplain [3].

Since the formulation of the flood-pulse concept and its extension [4,5], there have been substantial efforts to understand how river-floodplain biodiversity is influenced by hydrologic connectivity [6–9]. Such understanding would be key for the proper conservation and management of river-floodplains that are facing increasing pressure from human development and global climate change [10]. However, biodiversity-connectivity patterns appear complex and often inconsistent across different taxonomic groups. For instance, fish diversity generally peak in highly connected waterbodies [11,12], while amphibians showed the opposite patterns [5]. Benthic invertebrates and aquatic macrophytes displayed intermediate patters, with higher diversity often observed in dynamically connected waterbodies [6,13].

One of the main challenges in quantifying a general biodiversity response to hydrologic gradients in river-floodplains stems from the high spatio-temporal heterogeneity that characterise these ecosystems. For instance, only recently did studies specifically examined biodiversity changes through time and across different hydrologic stages [14–17], although conceptual models have long emphasised the stochastic nature of flooding events and the associated dynamics of floodplain habitats [4,5,18]. Partly, this is due to the challenge of monitoring rather unpredictable events in combination with pre- and post-disturbance data [e.g. 19].

In addition, the majority of studies have focused solely on the response of biodiversity and community metrics because these metrics are routinely quantified by monitoring programmes and used to guide ecological restoration [6,20]. However, biodiversity metrics, and their changes, ultimately result from the processes of community assembly [21]. Therefore, understanding how assembly mechanisms are influenced by flow variation and disturbance could provide additional insights into the dynamics of natural floodplains and their biodiversity. Research into community assembly has emphasised a distinction between processes driven by differences among species and their fitness (often called “niche”, or

“deterministic”) and processes that are mostly independent from these differences (“neutral” or “stochastic”). While the relative importance of these processes varies over a continuum [22], studies have illustrated how the relative effects of niche-based and stochastic processes depend on spatial scales, habitat heterogeneity, disturbance and ecological succession [23–26]. In particular, stochastic processes often appear to prevail in productive or benign habitats, while deterministic niche selection prevails under disturbed conditions [24]. However, disturbance can equally affect stochastic and deterministic processes depending on its intensity, timing, and duration [e.g. pulse *vs* press; 24]. Similarly, mature communities closer to equilibrium [sensu 27] appear regulated by local-scale deterministic processes while stochastic dispersal-based processes might be more evident in “young” or unsaturated communities [23,26]. It follows that any disturbance event able to reset communities to earlier stages will likely affect the relative importance of these assembly processes, but empirical assessment of temporal patterns is still scarce [16,28–30].

In this context, floodplains are ideal laboratories to test community assembly theory by examining how flow variations and local habitat conditions mediate the balance through time between stochastic and deterministic assembly processes. The patch-dynamics concept explicitly considered the frequent redistribution of organisms as a key driver in benthic community organisation within streams [31]. This dynamic perspective could equally describe biodiversity patterns in heterogeneous braided floodplains, but specific tests are mostly limited to regulated tropical rivers with predictable and gradual water level fluctuations [15,32,33]. However, temperate river floodplains are often characterised by unpredictable floods, which are expected to further increase in magnitude and frequency with climate change [34]. The effects of floods on the biodiversity of temperate floodplains still remains understudied [19,35]; at the same time, we are convinced that important insight could be gained by examining changes in apparent assembly processes during different hydrologic phases.

Here, we analysed a dataset on benthic Chironomidae and Oligochaeta at the genus and species level, sampled over an entire year from a meta-community of nine waterbodies across a lateral connectivity gradient in the semi-natural Tagliamento River (NE Italy; Fig.1). These taxa represent small-bodied organisms that are nearly ubiquitous along connectivity gradients in floodplains [6], have rapid generation times and are sensitive to flow variations [36] and can therefore be considered as ideal model organisms for testing community assembly theory [15]. Four bankful floods occurred during the study period (Fig. 2), allowing us to examine how hydrologic connectivity and flow variation determined spatio-temporal biodiversity patterns. Specifically we formulated the following questions:

q-i) How is the diversity of Chironomidae and Oligocheta communities influenced by the gradient of lateral hydrologic connectivity?
q-ii) Does hydrologic connectivity influence the response of communities to flooding events?
q-iii) To what extent do hydrologic connectivity and flood disturbance mediate the balance between deterministic and stochastic community assembly?

**Figure 1.**
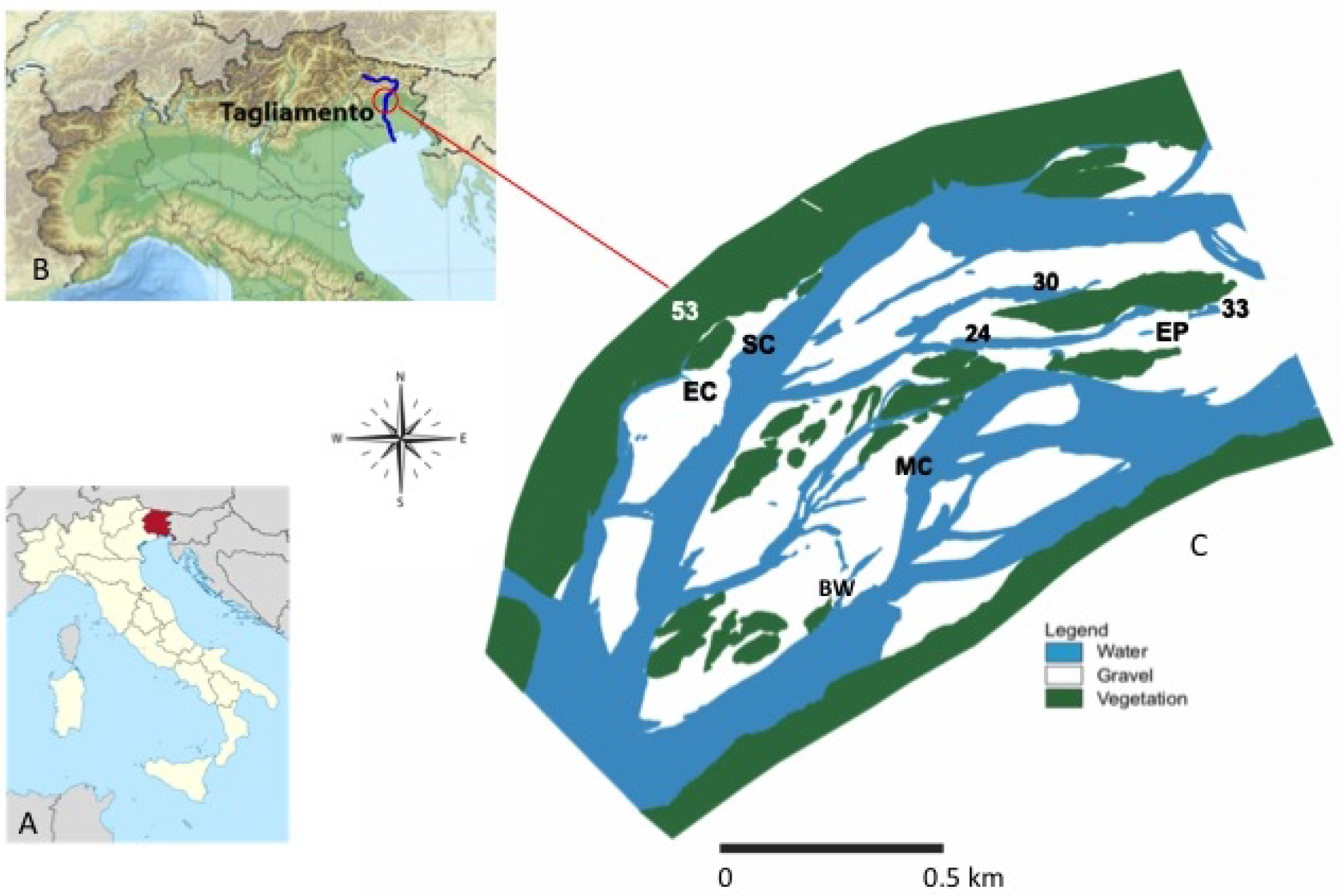
Map of the study area. (A) Map of the Trentino-Alto Adige region. (B) The Tagliamento river. (C) Detailed map of the study floodplain reach and the nine sampling sites (as in September 1999). MC = main channel; SC = side channel; EC = ephemeral channel; BW = backwater; EP = ephemeral pond.

**Figure 2.**
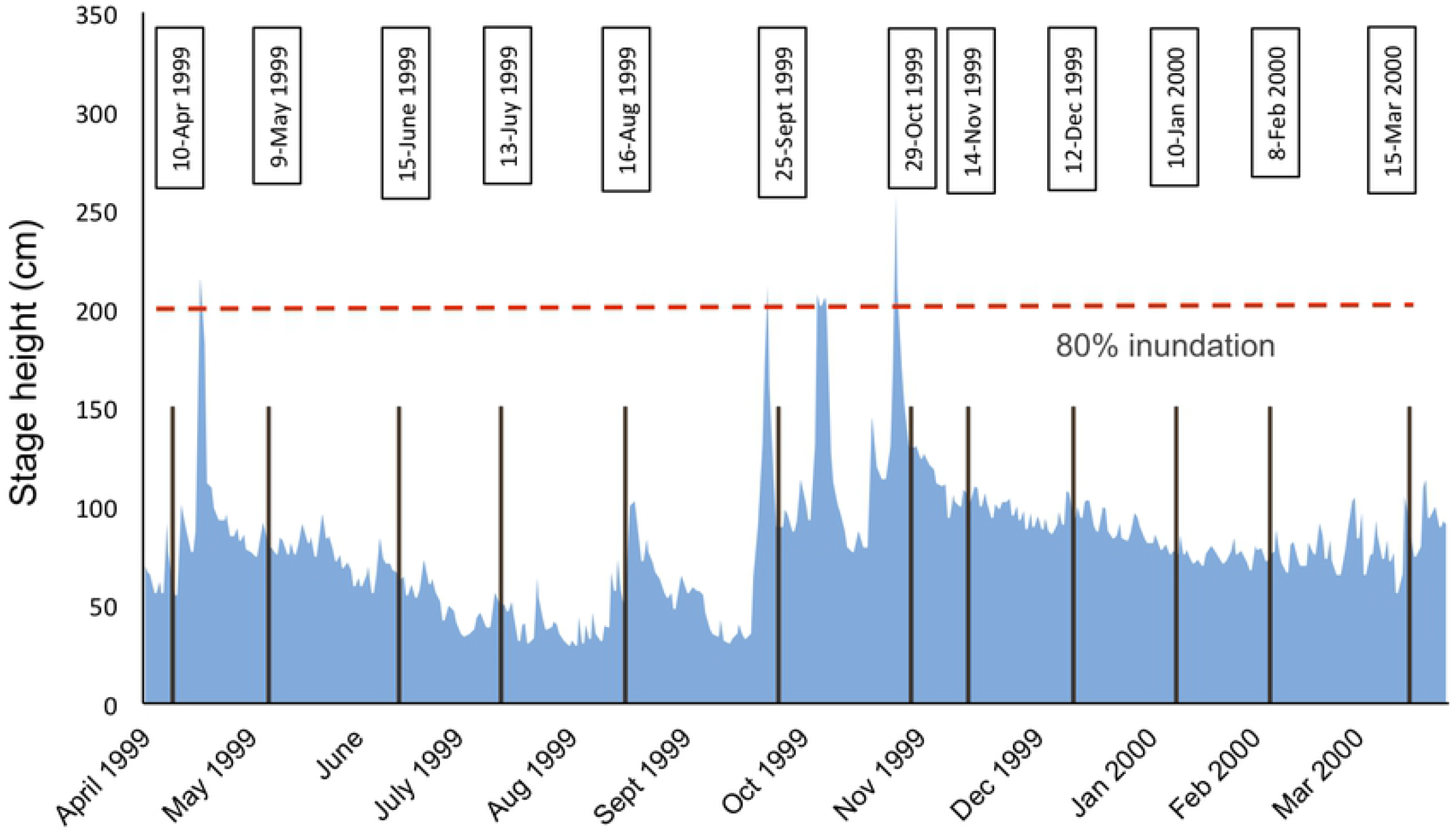
Hydrograph of the study period. Plot of the hydrograph showing stage height (cm) from April 1999 to March 2000, measured at the S. Pietro gauging station (3 km downstream of the study reach). Vertical bars indicate the time of invertebrate sampling with exact date listed in text boxes above. The red horizontal dashed line indicates the level where 80% of the study floodplain was inundated. Four main flooding events occurred during the sampling period.

Based on previous findings [6,16,20,37] and general community assembly theory [21,38], we were able to make specific predictions. First, we expected diversity to peak at intermediate levels of connectivity. Second, we expected disconnected waterbodies to be more severely affected by floods that those in the main and side channel, which should be more adapted to rapid flow variations. Third, the assembly of communities across the floodplain should proceed rather deterministically during stable hydrological periods (i.e. species-sorting prevailing); conversely, flooding events should increase the influence of stochastic processes related to random local extinction and re-colonisation processes.

## Methods

### Study area and design

The Tagliamento is a large gravel-bed river located in north-eastern Italy, in the Friuli-Venezia-Giulia region (Fig. 1A, B). It rises at 1195 m a.s.l. in the Carnian Alps and flows 170 km to the Adriatic Sea. The catchment covers 2580 km^2^ with more than 70% located in the Alps, and the average discharge is about 90 m^3^/s. High flows are caused by snowmelt (spring) and heavy rainfall (autumn) with discharge maxima of ~4000 m^3^/s [39]. The near pristine character of the Tagliamento is reflected in its complex channel morphology, dynamic flow and sediment regimes, and an idealised longitudinal sequence of constrained, braided and meandering sections [39].

The main study area is a 1 km^2^ island-braided reach in the middle section of the river corridor (river km 79.8 – 80.8; 135 m a.s.l.; Fig. 1C). The floodplain consists of channels, lentic water bodies (backwaters and ponds), gravel bars, vegetated islands, and the fringing riparian forest. The local climate has an Alpine character with a high precipitation of 2000 mm per year and a mean maximum air temperature of 17.6°C [40]. The waterbodies included in the study reflected a gradient of hydrologic connectivity with the main channel and included the main channel itself (MC), a secondary (SC) and one ephemeral (EC) channel, as well as one backwater (BW) with a downstream connection with the channel, and five ponds (named 24, 30, 33, 53 and EP) that differed in their location and distance from the channel. Water bodies 30 and 33 were embedded into the gravel matrix and located close and far from the channel, respectively. Water body 24 was associated with a vegetated island. Water body 53 was located at the edge of the riparian forest. The ephemeral pond (EP) was a bare-gravel pond. Ephemeral water bodies (EP and EC) dried in some periods during the investigation. The study was conducted at monthly intervals from April 1999 until March 2000, during which four bankful floods occurred (>200 cm stage height; Fig. 2), inundating c.80% of the active floodplain [41].

### Environmental data

Hydrological connectivity of each waterbody depends on the water level of the channel and the position of the specific waterbody relative to channel height. Here, using bi-weekly visits, connectivity was defined as the relative duration (% of time) that a waterbody was observed with a surface connection with the lotic channel.

Surface water temperature was recorded at hourly intervals using temperature data-loggers (VEMCO Minilog, Nova Scotia, Canada). Average daily temperature was calculated to characterize thermal heterogeneity across sites. Physicochemical variables were measured at monthly intervals from April 1999 until March 2000 prior to the sampling of benthic macroinvertebrates. However, data for the first and last month of the study were lost, so that only ten months of environmental data were included in the analyses. Variables included oxygen (O_2_; mg/l; Oxi 320, WTW, Germany), pH (pH 340, WTW, Germany), and specific conductance (μS cm^-1^, T_ref_ 20°C; LF 325, WTW, Germany). To minimize diel influences, all water bodies were sampled between 8:00 and 11:00 a.m. Surface water was collected in polyethylene bottles and kept cool (4°C) until analyses. Water was filtered through precombusted Whatman GF/F filters to separate particulate (suspended solids, particulate carbon, nitrogen and phosphorus) from dissolved components. Filters were frozen until analysis. All analyses were performed within four days of sample collection. Analytical methods for soluble reactive phosphorus (SRP), ammonia (NH_4_), nitrate (NO_3_), particulate nitrogen and phosphorus (PN, PP), total dissolved phosphorus (DP), dissolved nitrogen (DN), dissolved and particulate organic carbon (DOC, POC), and total inorganic carbon (TIC) were identical to Tockner et al. [42].

### Biotic data

Chironomidae and Oligochaeta were quantitatively sampled at monthly intervals from April 1999 until March 2000. In lotic water bodies, samples were collected by randomly placing a Hess sampler (415 cm^2^; mesh size: 100 μm) on the substrate and stirring the substrate with a metal rod for 10 sec to a depth of 5 cm (three replicates pooled per site and date). In standing water habitats, a diaphragm pump apparatus was used to collect benthic invertebrates from the surface within the enclosed Hess-sampler [43]. Ten litres were pumped from each Hess sample by holding the intake immediately above the sediment surface and by disturbing the surface with a rod for 10 sec to a depth of 5 cm (three replicates pooled per site and date). Samples were preserved in 4% formaldehyde. In the laboratory, all invertebrates were counted and sorted. Chironomidae and Oligochaeta were identified to the lowest taxonomic level, mostly genus and species (using identification keys reported in Appendix 1, in Supplementary Material). After removal of all invertebrates, coarse (>1.0mm) and fine (0.1-1.0mm) benthic organic matter (CBOM, FBOM, respectively) was dried and quantified as g/m^2^

### Data analysis

From the invertebrate data we calculated the observed taxonomic richness, Simpson diversity, as effective number of species of order q=2 [44,45] and density (individuals / m^2^). To represent temporal heterogeneity in local composition for each waterbody, we calculated the slope of the species accumulation curve based on random sampling over the twelve months period, limiting the effect of seasonality. Larger slopes reflect faster species turnover through time [e.g. 46]. Correspondence analysis (CA) based on log (x+1) densities was used to synthesise community composition for each waterbody and month.

To address question q-i, we used linear regressions to assess and plot the relationship between waterbodies hydrologic connectivity and the calculated community metrics, including richness, diversity, density, the slope of temporal species accumulation and mean composition (CA axes scores).

To address question q-ii, we first quantified, for each waterbody, the changes (in %) in community metrics (richness, density, composition) between each flooding event (i.e. between April-May, August-September and September-October). Then we regressed and plotted the changes in community metrics against hydrologic connectivity.

To address question q-iii, we used a set of complementary approaches to infer temporal changes in the dominant assembly processes. As recommended by Vellend (2010) we employed a multiple lines of evidence approach to evaluate changes in the signature of deterministic and stochastic processes through time. In the ecological literature, stochasticity conveys multiple concepts. Here, we followed Vellend’s (2010) notion in considering ‘neutral stochasticity’ as an assembly mechanisms being stochastic ‘with respect to species identity’. This is relevant here, as we must not confound environmental stochasticity (e.g. floods) with neutral stochasticity *(sensu* Vellend 2010).

First, following approaches from Chase and Meyers [47] and Kraft et al. [48], we examined whether beta-deviations (the departure of beta-diversity from what expected by the random distribution of the individuals) increased during stable hydrologic periods (reflecting increasing ecological determinism), and declined after flooding disturbance. Beta-deviations based on Bray-Curtis distances were derived from a quantitative null-model (999 simulations for each month) where every observed individual was randomly placed in a waterbody within the metacommunity until every individual was placed. This maintained the overall metacommunity abundance distribution and richness, but randomised the individuals’ location and the spatial aggregation within and among species [49]. Beta-deviations thus control for changes in local richness and also inform on potential assembly mechanisms as the distance of the observed deviations from the null-expectation can indicate the relative influence of deterministic assembly processes as opposed to more stochastic ones [47,48].

Second, following approaches from Jenkins [23], Ellwood et al. [50] and Larsen and Ormerod [26], we assessed how flow disturbance influenced species co-occurrence patterns. Specifically, we examined whether stable flows allowed the formation of non-random cooccurrence patterns across the floodplain habitats (reflecting species-sorting driven by biotic interactions and environmental filtering), and whether flooding events shifted the patterns towards random. Species co-occurrence was calculated from presence-absence data using the C-score index [51], which is the average number of all checkerboard units (mutual exclusion) calculated for each species pair in the metacommunity. Although the ability of the C-score index to effectively discriminate segregation and aggregation has been discussed [52], its use in combination with the fixed-fixed null-model algorithm has good power to identify nonrandom patterns, shows low Type I and Type II error rates and is less affected by variations in species richness compared to other indices [53]. This was ideal for our analyses as we were interested in assessing how the formation of non-random structure changed through time while controlling for concomitant variations in local richness. The C-score was expressed as standardised effect size (SES) for each month as:

SES = (observed C-score – mean simulated C-score) / SD simulated C-scores.

If this index was larger or smaller that what was expected by a random distribution of individuals, it indicates that there is respectively less (species segregation) or more (aggregation) co-occurrence than expected by chance. Non-random patterns were detected using the fixed-fixed null-model [54] in which local site richness and species frequencies in the null metacommunities were equal to those originally observed.

Finally, we examined whether the influence of local habitat features was greater during stable flow periods (as species are sorted according to their niche requirements), compared to periods after flooding events, which were expected to weaken the match between local environmental conditions and species composition. We thus used redundancy analysis to quantify the variance in community composition explained by local habitat features each month (expressed as adjusted R-square), and included the available physico-chemical data and degree of hydrologic connectivity as explanatory tables, and the species log (x+1) density data as response table.

All analyses and plots were conducted in R [55] using the Vegan package and null-model codes provided by Tucker et al. [56].

Data associated with this work are available from the Dryad repository.

## Results

### Abiotic patterns

The main physicochemical characteristics of the study sites are reported in Table S1 (Supplementary Material, SM). Mean values of SRP (R^2^=0.39; p=0.057), DP (R^2^=0.6; p=0.014), and POC (R^2^=0.44; p=0.04) increased with hydrologic connectivity, although the relationship was only marginally significant for SRP. Similarly, a marginally significant increase in oxygen concentration (p=0.09) and pH (p=0.06) with connectivity was observed.

### Biodiversity-connectivity patterns (question q-i)

A total of 82 Chironomidae and 34 Oligochaeta taxa were collected during the study. Chironomidae were the most abundant. In particular, the sub-genus *Euorthocladius sp.* was the most abundant taxon, with a mean density (all sites and months) of 1983 individuals / m^2^, followed by *Micropsectra sp.* (490 individuals / m^2^). The most widespread taxa (both Chironomidae) were *Polypedilum sp.* and *Tanytarsus sp.,* which were observed in eight out of the nine waterbodies sampled. Overall mean density varied greatly across floodplain habitats and was an order-of-magnitude higher in the main-channel, side-channel and backwater (10118, 10110, and 6396 ind / m^2^, respectively; Table S1), compared to the more isolated waterbodies (range: 90 – 690 ind / m^2^). Mean observed species richness showed a positive relationship with the degree of lateral connectivity (Fig. 3A; R^2^= 0.5; p=0.04), whereas no relation was observed for mean diversity of order q=2 (Fig. 3B). Although the highest richness values were observed at around 75% connectivity, there was no support for a nonlinear relationship and thus no evidence for a peak in mean richness at intermediate levels of connectivity. Mean density showed a rather strong positive relationship with connectivity (Fig. 3C; R^2^=0.9; p<0.001). Overall, richness-connectivity relationships were variable through time and exhibited both linear and quadratic patterns, depending on sampling date (SM; Fig.S1).

**Figure 3.**
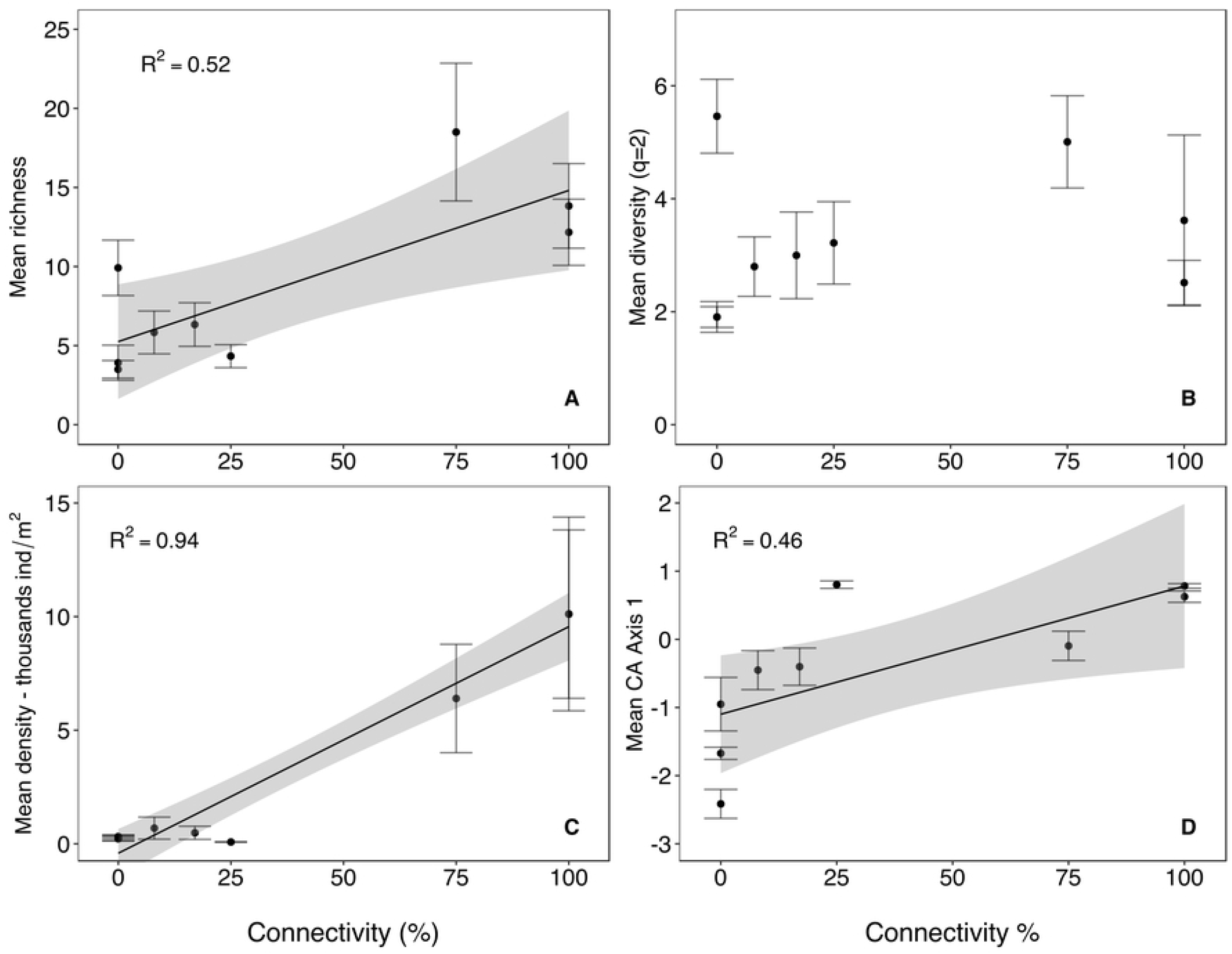
Biodiversity – connectivity relationships. Relationship between mean (±SE) taxonomic richness (A), mean Simpson diversity of order q=2 (B), mean density (C) and mean CA (Correspondence Analysis) axis 1(D) with lateral hydrologic connectivity across nine waterbodies in the Tagliamento floodplain (NE Italy). Grey area = 0.95 confidence interval of regression.

Taxonomic composition was influenced by connectivity too, with mean CA axis 1 (explaining 11% of the overall variance in taxonomic composition across sites and months) increasing linearly along the connectivity gradient (Fig. 3D; R^2^=0.46). In addition, temporal variation in taxonomic composition was higher in disconnected sites, indicating higher temporal species turnover rates with decreasing connectivity (SM; Fig.S2). This is evident from the negative relationship between the rate of temporal species accumulation, calculated for each site, and the connectivity gradient (Fig.4; R^2^=0.4).

**Figure 4.**
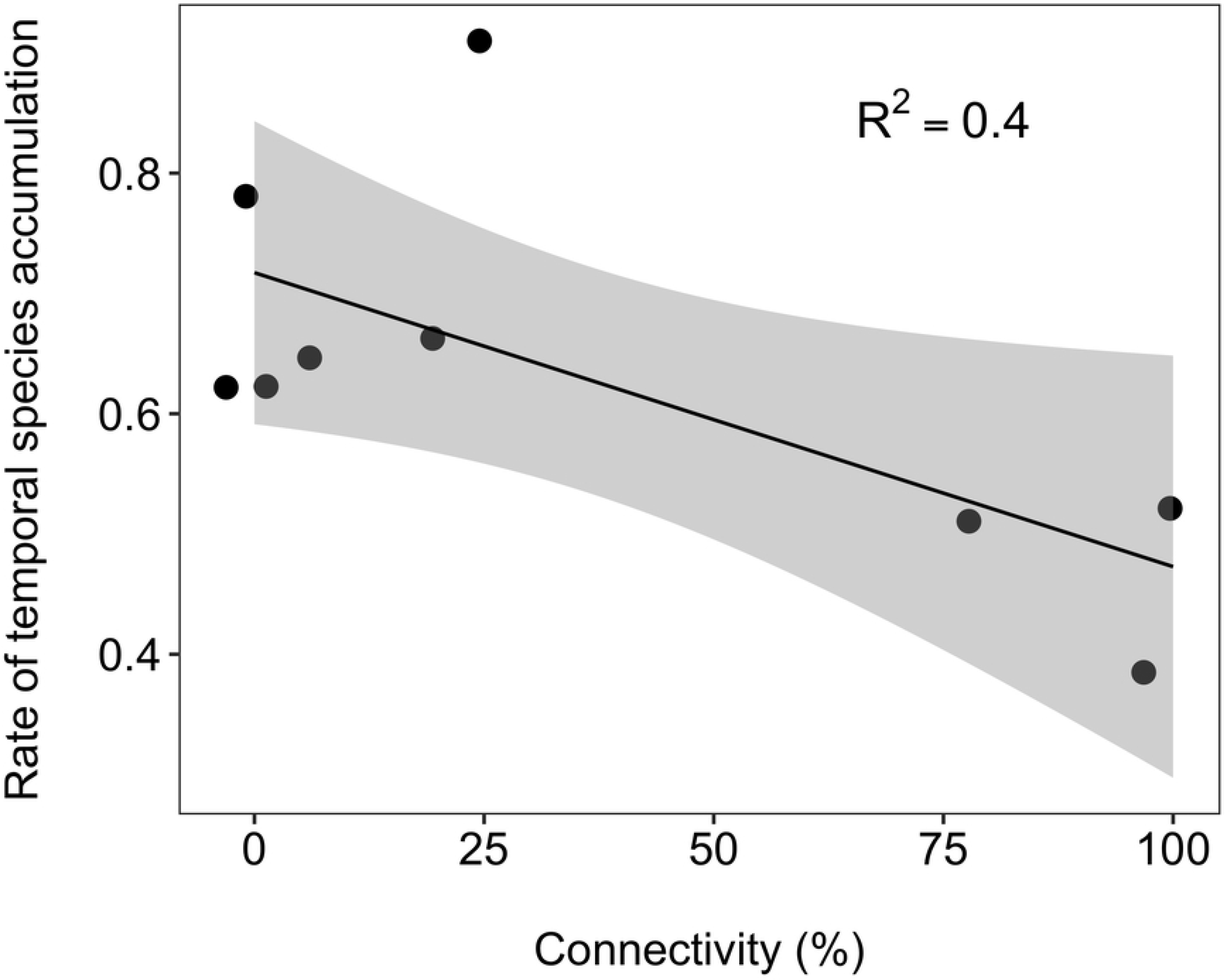
Temporal species accumulation. Relationship (±0.95 CI) between the rate of temporal species accumulation and lateral hydrologic connectivity across nine waterbodies in the Tagliamento floodplain (NE Italy)

### Habitat-specific response to floods (question q-ii)

Changes in taxonomic richness, density and community composition, in response to flooding events, were not related to the connectivity gradient, which was in contrast to our predictions. Fig. 5 shows that the percentage changes in richness and density between consecutive months, separated by flooding events (i.e. months: Apr-May, Aug-Sept and Sept-Oct), were unrelated to the degree of hydrologic connectivity. Similarly, changes in community composition after flooding, as reflected by changes in the first CA axis, were also unrelated to connectivity (SM Fig.S3).

**Figure 5.**
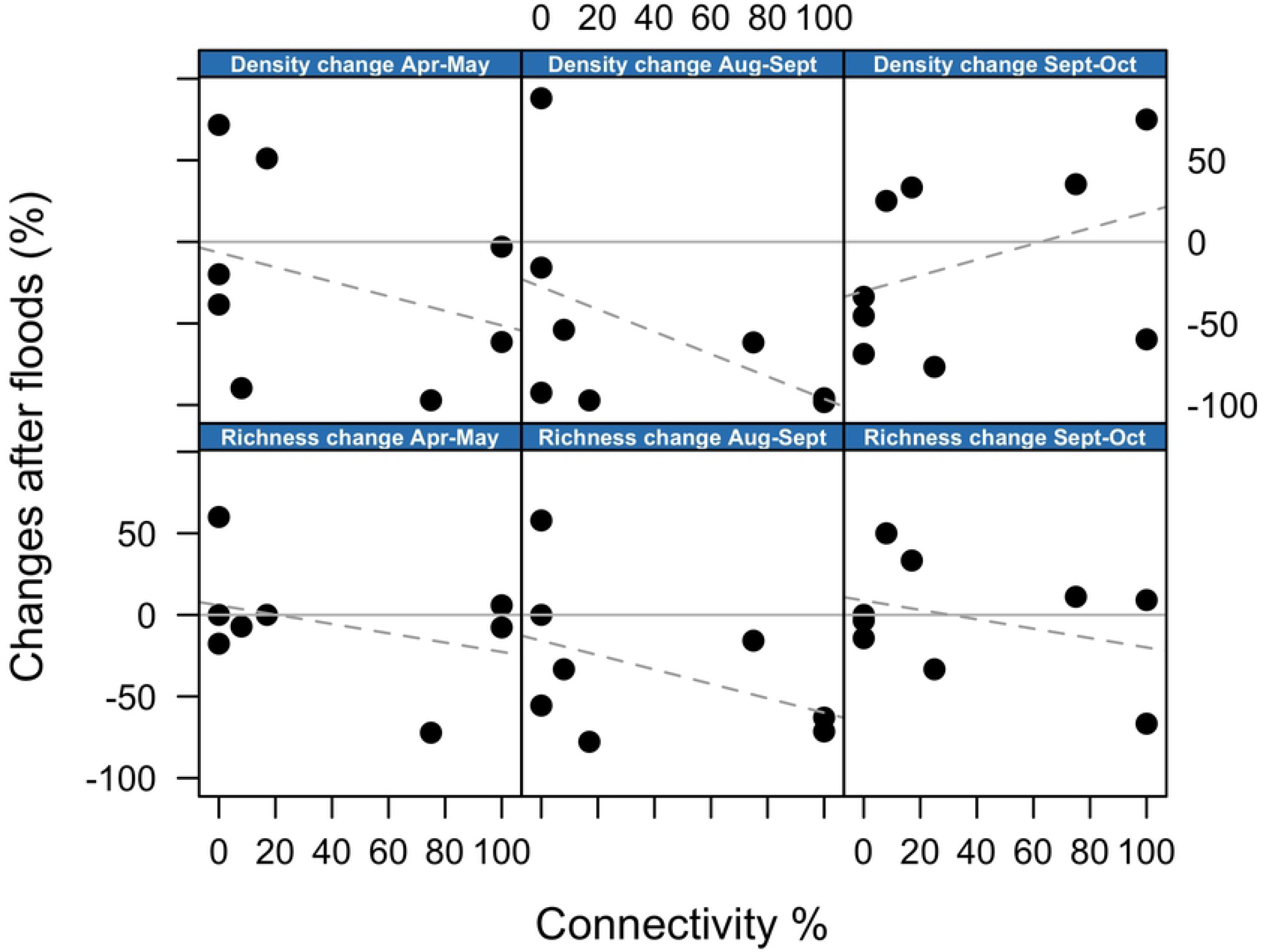
Change in community metrics after floods. Relationship between hydrologic connectivity and percentage changes in density and richness after each flooding event. Each panel shows changes in either density or richness between consecutive months during which flooding occurred (e.g. April-May). Zero change is indicated by a solid grey horizontal line. No significant correlations were observed. For visualisation purposes, a dashed grey line indicates the direction of the relationship.

### Community assembly mechanisms (question q-iii)

We used multiple independent approaches to assess temporal changes in apparent assembly mechanisms. Fig. 6 shows the temporal changes in taxonomic composition as reflected by each sites’ CA axis 1 scores. Flooding events (represented by blue vertical lines) appeared to cause a faunal convergence across the different floodplain habitats (i.e. converging towards the main and side channel), whereas months of stable flows (e.g. months 5 to 8) were characterised by a progressive faunal divergence of the habitats, which partly reflected the gradient of hydrologic connectivity.

**Figure 6.**
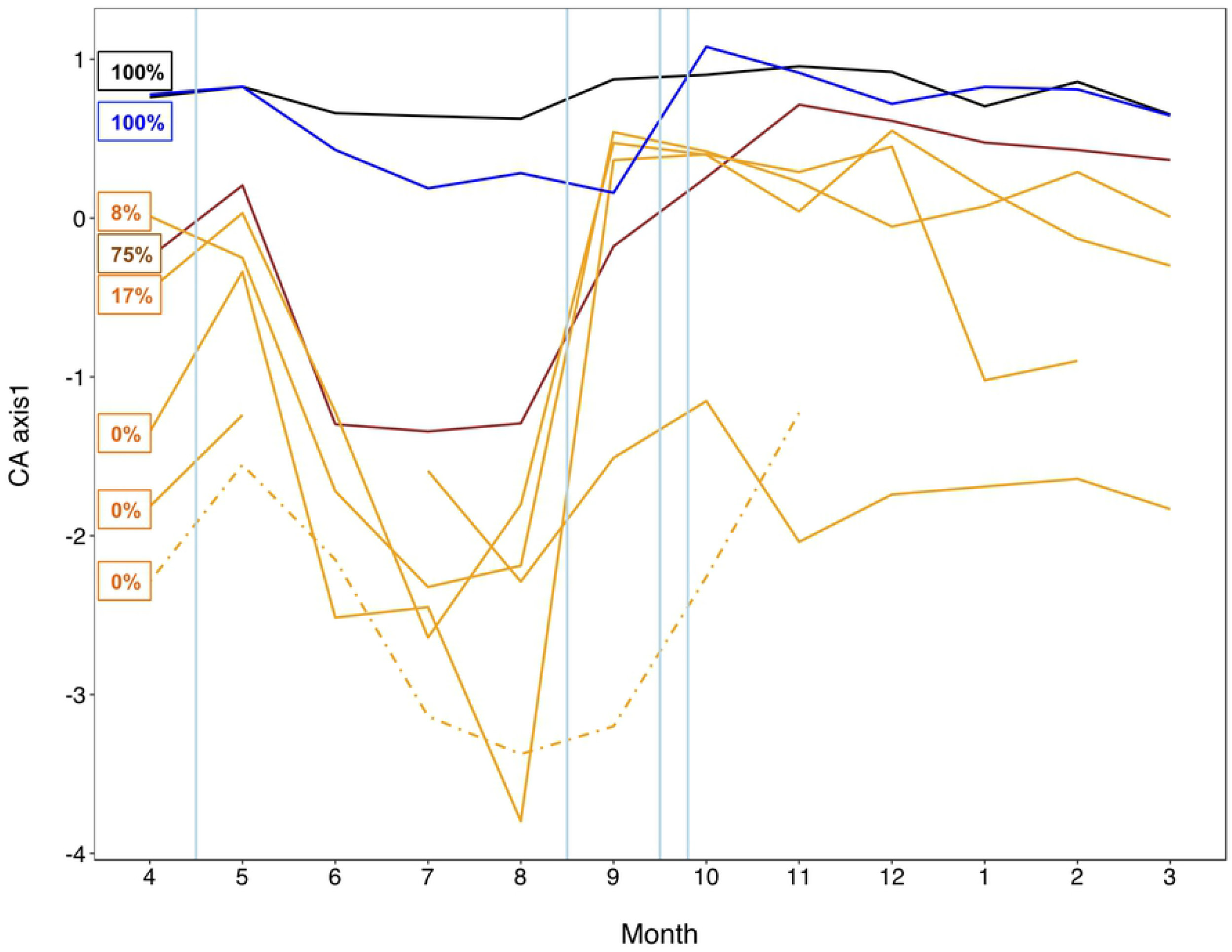
Temporal changes in community composition. Correspondence Analysis (CA) axis scores for each floodplain waterbody over the hydrologic year. The colour code follows the gradient of lateral hydrologic connectivity (%) indicated in the text boxes. Black and blue lines represent the main and side channel, respectively. The brown line indicates backwater, while the others, mostly isolated, waterbodies are shown in orange. The dashed line indicates the ephemeral pond. The ephemeral channel, inundated for only three months, is not shown. Vertical blue lines indicate the occurrence of bankful floods. Temporal patterns show that the composition of the floodplain waterbodies tended to converge (towards a ‘main channel’ condition) after the floods, but to diverge during months of stable hydrologic conditions, reflecting differential species sorting.

Null model analyses provided further insights. In particular, mean beta-deviation appeared to decline after flooding events and to increase steadily during stable flow conditions, reflecting progressive species sorting across the floodplain habitats (Fig. 7A). Patterns in co-occurrence were also consistent with a stronger effect of species sorting during stable flows. Species cooccurrence appeared random (z-score < 1.96), or independent, after flooding events and progressively shifted to non-random after 2-3 months of stable flows (Fig. 7B). Finally, the proportion of variation in taxonomic composition explained by local environmental variables ranged between 0.07 – 0.40 during stable flows, and between 0.01 – 0.08 after floods (Fig. 7C; note that data from the first and last month of study were lost). This suggests that, on average, taxonomic composition was less predictable after flooding events.

**Figure 7.**
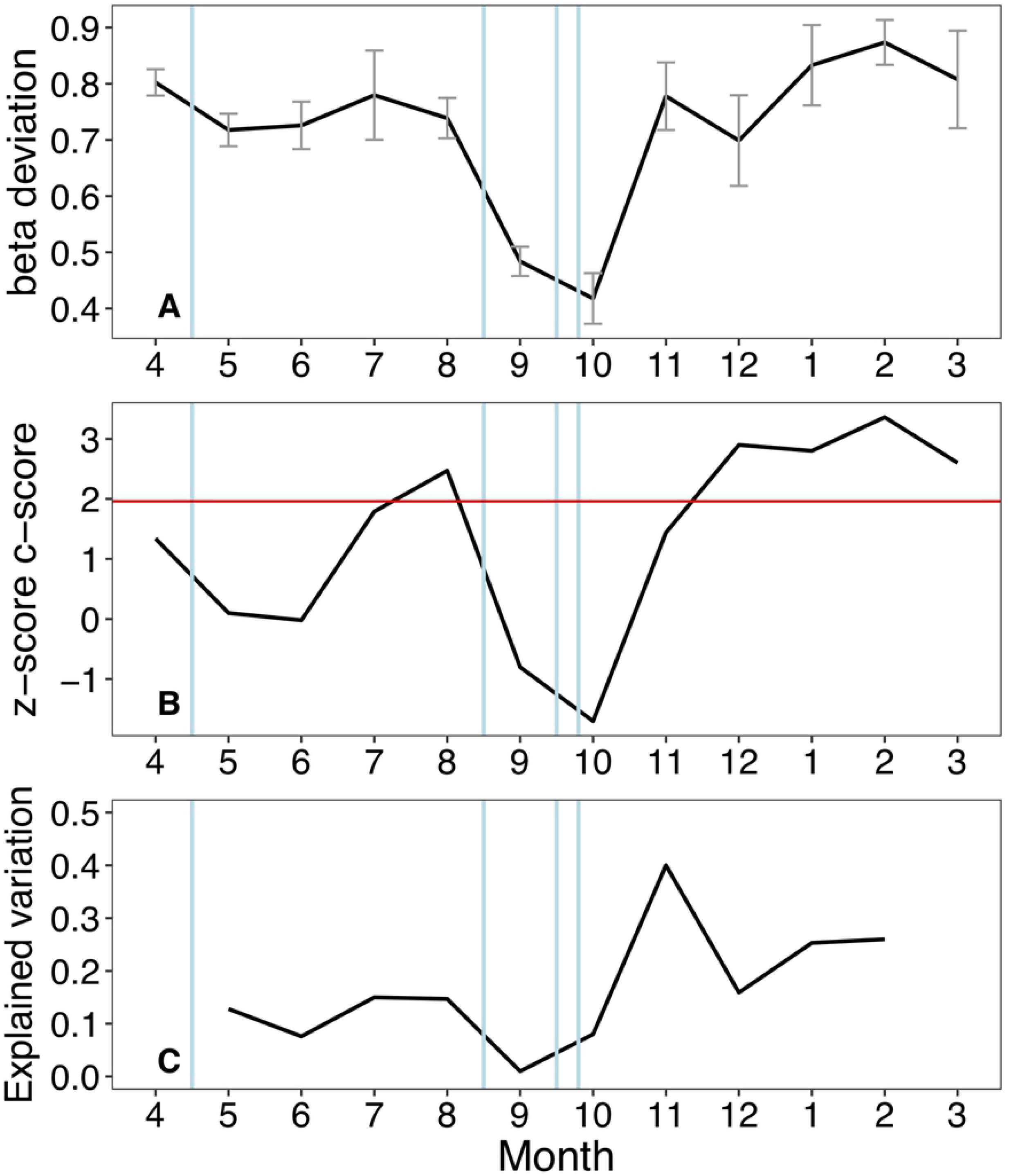
Inferring community assembly mechanisms. Temporal patterns in mean pairwise beta-deviation across waterbodies (A), species co-occurrence z-scores (B) and the proportion of variation in composition explained by local environmental parameters (C). Vertical blue lines indicate the occurrence of bankful flood events. The three indicators consistently showed that flooding weakened the signal of deterministic assembly processes. In (A), mean beta-deviation declined after each flood. In (B), months with stable hydrologic conditions were associated with non-random species co-occurrence patterns (above the red line at z-score = 1.96), which shifted to random patterns after flooding. In (C), flooding tended to decrease the match between local abiotic factors and species composition. Abiotic data for the first and last months were not available.

These temporal changes in the apparent assembly mechanisms occurred despite no evidence for a homogenising effect of floods on the abiotic characteristics of the floodplain sites. Indeed, mean Euclidean distance among floodplain waterbodies, according to their physicochemical parameters, did not decline consistently after flooding events (SM. Fig S4).

## Discussion

This is among the few studies that assessed biodiversity dynamics and assembly mechanisms in active river floodplains over an entire hydrologic year [e.g. 14-17]. Results indicated that in the Tagliamento floodplain, invertebrate richness, diversity and density manifested complex and temporally variable relations with hydrological connectivity. Disconnected waterbodies were characterised by faster rates of intra-annual turnover in faunal composition. However, and contrary to expectations, connectivity did not appear to influence the magnitude of change in community metrics and composition after flood events. An additional key finding is that hydrologic connectivity and flow variation, single and in concert, regulated the dynamic balance between deterministic and stochastic community assembly, highlighting the importance of employing a temporal perspective in the study of assembly mechanisms, especially in dynamic systems such as river floodplains [17,19,56].

### Abiotic patterns

Hydrological connectivity controls a range of local environmental conditions in floodplain habitats [6]. In the braided floodplain of the Tagliamento River, the concentration of phosphorous and particulate organic carbon was higher in connected waterbodies – as expected, reflecting direct inputs from the main channel. This is in agreement with other studies reporting higher nutrient concentration in hydrologically connected sites [6,20]. Disconnected waterbodies also tended to be more acidic and have lower oxygen concentration than connected sites, as also observed in other active floodplains [16,57] However, the influence of local environmental parameters on resident communities changed over time and generally declined after flooding events.

### Biodiversity-connectivity patterns

The relationship between lateral connectivity and floodplain biodiversity has received considerable attention [6,12,43], but few consistent patterns have emerged so far. The intermediate disturbance hypothesis (IDH) has often guided predictions, and some support has emerged with higher invertebrate diversity often observed in waterbodies that connect with intermediate frequency [3,5,20]. In our study system, connected waterbodies supported much higher densities of invertebrates, which determined a linear relationship between richness and lateral connectivity. However, richness-connectivity relations were variable through time with both linear and hump-shaped patterns evident in different months. Overall, the present results provided no support for the IDH, in part because our study sites were not evenly distributed over the connectivity gradient (i.e. no data at intermediate connectivity level).

The Simpson diversity (q=2), which is influenced by the most abundant taxa, was not related to hydrologic connectivity. This indicates that the higher richness observed in connected waterbodies was accompanied by uneven species abundance distribution and was partly driven by a sampling effect due to higher local densities [44]. Biodiversity-connectivity patterns appeared thus highly variable as also observed previously [3,58]. However, direct comparisons with other studies are difficult because our analyses focused on Chironomidae and Oligochaeta, two of the most diverse and ubiquitous invertebrate groups in floodplains. However, both groups are often underrepresented in benthic invertebrate studies [6]. Sporka and Nagy [36] reported a decline of Oligochaeta and Chironomidae density with a transition from lentic to lotic conditions in side channels of the Danube River (Slovakia). Similarly, recent work from the Parana’ floodplain [59] recorded the highest density of chironomids in completely disconnected waterbodies, a result apparently in contrast with ours. These findings, together with our own, further highlight that the influence of hydrologic connectivity on floodplain biodiversity cannot be reduced to a single gradient [3,8]. Including a temporal dimension can provide additional insight into these cases. In the Tagliamento floodplain, disconnected waterbodies were characterised by faster rates of species turnover through time (i.e. faster rates of temporal species accumulation), reflecting a greater variability compared to permanently connected sites. In apparent contrast, Starr et al [16] reported higher macroinvertebrates turnover in lotic compared to lentic habitats, but this was attributed to the strong environmental filtering associated with widespread anoxia in the more isolated habitats. The flood pulse concept [4] originally predicted higher biodiversity in dynamic floodplain waterbodies compared to the main channel, but specific temporal investigations are still scarce. The disproportionate contribution of lateral habitats to regional diversity has been demonstrated by extensive surveys across different floodplains [8,9], and our results suggest that the contribution of lateral habitats to total floodplain diversity increases even more when the temporal dimension is included.

### Habitat-specific response to floods

In the Tagliamento floodplain, hydrologic connectivity did not influence the response of Chironomidae and Oligochaeta community metrics to flood events. This was unexpected since we predicted flooding to have a stronger effect on lateral disconnected waterbodies compared to the main channel, where organisms should be adapted to rapid flow variations [4,5]. Although none of the trends were significant, Fig. 5 suggests that flooding was actually associated with slight reductions in richness and density in the most connected waterbodies (except for density change between Sept-Oct). This suggests that, although communities in the main channel might be adapted to flow variation, the disturbance associated with sediment scouring and high water velocity during floods was stronger in the main channel compared to lateral habitats.

### Community assembly mechanisms

The general scheme of stochastic-deterministic processes has guided most contemporary assembly studies, with consensus emerging about a continuum perspective, where both processes operate simultaneously [22]. However, key questions remain about the influence of environmental change on the specific outcomes. For instance, the influence of disturbance or environmental harshness on assembly processes appeared contingent on the type of disturbance, its duration as well as the spatial scale considered [24,25,60,61]. Similarly, ecological succession also influences the relative importance of assembly mechanisms, with deterministic species sorting and biotic interactions prevailing in mature communities [23], as opposed to ‘younger’ communities regulated by more stochastic processes of dispersal and priority effects [29,62]. Predictability and magnitude of environmental change can thus affect our ability to infer assembly mechanisms. Therefore, including the temporal dimension in the study of community assembly can provide important insight, and dynamic river-floodplains represent ideal natural laboratories to advance our understanding of assembly mechanisms in a changing world [16,17,30].

Using a suite of complementary approaches, our results were consistent with the hypothesis that, in the Tagliamento floodplain, floods represented a key factor that weakened the signal of deterministic species sorting and apparently re-set communities to earlier successional stages. Trajectories of community dynamics diverged across floodplain habitats during stable hydrologic periods, but converged again towards a ‘main-channel’ condition after the floods. Results further indicated that the effects of niche-driven species sorting and potential biotic interactions were stronger during stable hydrologic periods, when the metacommunity exhibited higher beta-deviations, non-random species co-occurrences and taxonomic composition that, on average, better matched the local environment. These patterns changed consistently in post-flood periods where neutral stochasticity became more apparent [sensu 38]. Such dynamics are in agreement with community assembly theory [38,63] and with river-floodplain models such as the patch-dynamic [31], flood pulse [4] and river ecosystem synthesis [18] models, which clearly emphasised the dynamic balance between stochastic and deterministic forces in regulating riverine communities. Empirical tests of dynamic assembly mechanisms lagged behind these conceptual models, but evidence from floodplain systems is growing. Arrington et al [32] were among the first to use contemporary assembly approaches to relate the timing of flood disturbance with the formation of non-random co-occurrences in fish communities in tropical floodplains. Similar findings were later reported also by Fernandes et al. [33]. Co-occurrence patterns have been traditionally analysed to infer assembly processes related to species interactions [64], but non-random co-occurrences likely reflect the combined effects of biotic interactions and deterministic habitats filtering [50,65]. Previous experiments demonstrated how ecological succession could alter species cooccurrence in predictable ways with random patterns prevailing in disturbed and ‘early-stage’ metacommunities [23,26,50]. In agreement with Arrington et al. [32] and Fernandes et al. [33], flooding in the Tagliamento was consistently associated with the shift from non-random to random species associations supporting the hypothesis that communities were reset to earlier successional stages. The exact mechanisms involved cannot be identified in observational studies such as this, but a weakening of biotic interactions, organisms’ redistribution, and a decrease in habitat heterogeneity are all possible. A decline in floodplain heterogeneity during floods has been proposed as one of the few ‘general principles’ of floodplain ecology [37]. Surprisingly, we did not observe such homogenising effect in the measured abiotic parameters across the waterbodies. This could be attributed to the fact that, for practical reasons, water samples were not collected during the peak of the floods, but always few days later. Therefore, it is possible that we were not able to fully quantify the short-term responses of physico-chemical parameters that likely varied rapidly during flood events. Alternatively, it is possible that we did not measure some important abiotic factors, such as local substratum characters, which could potentially be homogenised across the floodplain during floods.

Conversely, temporal patterns in biotic homogenisation, as expressed by changes in mean beta-deviations, appeared to clearly match the occurrence of the floods. It is thus possible that the time lag after which any homogenising effects of floods may become apparent differs between the biotic and abiotic component [37,66]. While declining beta-diversity with increasing floodplain water levels has been observed across a range of taxonomic groups [66,67], and used to infer assembly mechanisms, quantifications of null-based beta deviations are much scarcer [17,68]. Raw beta-diversity values can be influenced by stochastic variations in local and regional diversity, making it difficult to disentangle process-based changes from those resulting from chance variations in local composition and richness [47]. Beta deviation values that control for such stochastic variation are therefore particularly useful to quantify temporal changes in dynamic systems such as floodplains, with larger deviations reflecting deterministic assembly processes related to species sorting and competitive biotic interactions [48,49]. In agreement with patterns in species co-occurrence, flooding in the Tagliamento appeared to temporary weaken the importance of these deterministic processes, reflecting the increased likelihood of dispersal among locations and the decline in beta-deviations. While these findings are in line with the hypothesis that floods promoted biotic homogenisation [37,67], other studies did not observe a relationship between floodplain beta-diversity and hydrologic variations [68,69], and suggested that antecedent conditions related to regional species pool and local population density represented buffering factors (ecological memory) that maintained stable local alpha- and beta-diversity. However, these studies were all set in larger tropical floodplains characterised by rather predictable and gradual flood pulses. In the Tagliamento floodplain, hydrologic variations occurred in both spring and autumn and were relatively rapid thus representing a more severe disturbance.

Besides null-model based assessments, assembly processes have been inferred by partitioning the variation in community composition into environmental and spatial components [70], whose relative importance should reflect niche-based and more stochastic dispersal-based processes, respectively. For instance, this approach has been used to conclude that invertebrate metacommunities in floodplain ponds were mostly structured by niche processes [71], or a combination of dispersal and niche processes [72,73]. However, snapshot observations can be misleading because the relative influence of environmental and spatial drivers can change through time, especially in ecosystems frequently influenced by hydrological disturbance [28,74]. Our results provide support for this dynamic view; using the same set of environmental parameters (including the degree of connectivity), we showed that flooding affected the extent to which communities matched local abiotic conditions (i.e. the predictive power of the environmental component). Although data were not available for the first and last month in this case (i.e. the effect of the first flood cold not be assessed), these patterns were consistent with those of beta-deviation and co-occurrence in indicating that flooding tended to decrease ecological determinism across the floodplain habitats. However, relatively large changes in explained variation occurred also during months of stable flows, indicating that other factors besides flooding influenced the match between local abiotic conditions and resident communities.

## Conclusion

Understanding how fluvial processes and connectivity shape alpha and beta diversity patterns has been central in the study of floodplain ecology [8,9,11]. Along with other recent investigations, this work highlights the importance of the temporal scale in the study of such dynamic ecosystems [16,17,74]. The Tagliamento River represents the last morphologically intact river in the Alps and can be considered as a model ecosystem for large temperate rivers [40]. In the braided section of the river, flow variation is a dynamic factor that not only regulates local invertebrate abundance and richness, but also changes the main assembly processes across the floodplain. While these findings are in agreement with general models of river-floodplain ecology and community assembly theory, unequivocally identifying the underlying assembly mechanisms is not possible from patterns derived from observational data. This is a common issue in most community assembly investigations unless experimental manipulations are involved [26,63]. Ideally, repeating this study in the absence of floods or with different flood timing would provide further support to our findings independently from any seasonal or phonological influence. However, floods occurred in both spring and autumn with patterns that were consistently observed from multiple approaches.

Overall, the present results further highlighted the vital role of hydrologic connectivity and variable flow regimes in supporting lateral floodplain habitats, which are hotspot of biodiversity [8]. These are the landscape elements that disappear first as a consequence of channelization and flow regulation, with 90% of former floodplains in Europe already lost or degraded [75]. Successful management and conservation of floodplains depends on the thorough understanding of their spatio-temporal dynamics [75]. However, the wider implications of the present findings would benefit from a deeper understanding of the links between community assembly and ecosystem functioning [76]. In fact, while the link between biodiversity and ecosystem functioning is well established [e.g. 77], the role of different assembly processes, which determine the current biodiversity in the first place, is mostly unknown. However, changes in biodiversity deriving from either stochastic or deterministic processes can affect ecosystem functioning in markedly different ways depending on the functional traits of the species involved. Research in this direction is at its infancy [78] and natural floodplains could serve as ideal model systems for developing and testing new hypotheses.

## Acknowledgements

This research was supported by ETH Forschungskommission grant No. 0-20572-98. S. Larsen was supported by funding from the European Union’s Horizon 2020 Research and Innovation Programme under grant agreement No. 748969 with an Individual MSC Fellowship.

The authors also thank Luana Botinelli, James Ward, Dimitry van der Nat and David Arscott for help in the field.

## Supplementary Material – Figure & Table Caption

**Appendix 1** – Identification keys used in the study

**Table S1. Main characterisitcs of the study floodplain**

Degree of hydrologic connectivity and main physico-chemical characteristics of each floodplain waterbody. Density and richness of the invertebrate communities is also shown.

**Fig. S1. Richness-connectivity over time**

Correlation patterns between observed richness and lateral hydrologic connectivity (%) for each month. A smooth line is included to show the variation in the pattern through time.

**Fig.S2. Temporal change in faunal composition**

Temporal variation in the composition of each floodplain waterbody as expressed by the first Correspondence Analysis (CA) factorial plane. For each waterbody, the degree of hydrologic connectivity (%) is also shown. Each point represent one month. Decreasing variation (especially along the first axis) with increasing connectivity is evident.

**Fig. S3. Composition changes after floods**

Relationship between hydrologic connectivity and changes in taxonomic composition as expressed by the first Correpondence Analysis (CA) axis scores. Each panel shows changes in CA axis 1 scores between consecutive months during which flooding occurred. No significant correlations were observed, but for clarity a dashed line was included to show the direction of change.

**Fig.S4. Temporal changes in abiotic distance between sites**

Temporal changes in mean (SE) Euclidean distance among floodplain waterbodies based on the main physico-chemical parameters. Blue vertical lines represent flooding events. There is no consistent indication of a decline in distance (i.e. abiotic homogenisation) with the occurrence of floods. Data from the first and last months (month 4 and 3, respectively) were not available.

